# Fear extinction is regulated by long noncoding RNA activity at the synapse

**DOI:** 10.1101/2022.03.30.486308

**Authors:** Wei-Siang Liau, Qiongyi Zhao, Adekunle Bademosi, Rachel Gormal, Hao Gong, Paul R. Marshall, Ambika Periyakaruppiah, Sachithrani U. Madugalle, Esmi L. Zajackowski, Laura J. Leighton, Haobin Ren, Mason Musgrove, Joshua Davies, Simone Rauch, Chuan He, Bryan C. Dickinson, Xiang Li, Wei Wei, Frédéric A. Meunier, Sandra M. Fernandez Moya, Michael A. Kiebler, Bharath Srinivasan, Sourav Banerjee, Michael Clark, Robert C. Spitale, Timothy W. Bredy

## Abstract

Long noncoding RNAs (lncRNAs) represent a multidimensional class of regulatory molecules involved in many aspects of brain function. Emerging evidence indicates that lncRNAs are expressed at the synapse; however, a direct role for their activity in this subcellular compartment in memory formation has yet to be demonstrated. Using lncRNA capture-seq on synaptosomes, we identified a significant number of lncRNAs that accumulate at synapses within the infralimbic prefrontal cortex of adult male C57/Bl6 mice. Among these is a splice variant related to the stress-associated lncRNA, Gas5. RNA immunoprecipitation followed by mass spectrometry and single molecule imaging revealed that this Gas5 isoform, in association with the RNA binding proteins G3bp2 and Caprin1, regulates the activity-dependent trafficking and clustering of RNA granules in dendrites. In addition, we found that cell-type-specific, state-dependent, and synapse-specific knockdown of the Gas5 variant led to impaired fear extinction memory. These findings identify a new mechanism of fear extinction that involves the dynamic interaction between local lncRNA activity and the coordination of RNA condensates in the synaptic compartment.

## Introduction

The extinction of conditioned fear is a critically important adaptive behaviour that is driven by activity in the infralimbic prefrontal cortex (ILPFC). Like other forms of learning, long-lasting memory for fear extinction depends on coordinated changes in gene expression ^1-3^. Although significant progress has been made in revealing the mechanisms that regulate this process, a complete understanding of the molecular code underlying the formation of fear extinction memory is still lacking. Long noncoding RNAs (lncRNAs) have emerged as key regulatory molecules associated with a variety of important biological processes, including gene regulation, translation, and RNA trafficking ^4-7^. Central to their multifunctional capacity, lncRNAs are expressed in a cell-type-specific and spatiotemporal manner and are highly enriched in the brain. Several lncRNAs have been found to accumulate in the synaptic compartment in response to neural activity. For example, the lncRNA BC1 appears to facilitate local translation through an interaction with the ribosome and translation initiation factors ^7-9^, while ADEPTR accumulates in dendrites where it mediates activity-dependent changes in synaptic plasticity ^10, 11^ suggesting a potential role for the localised expression of lncRNAs in the regulation of synaptic processes underlying memory.

Previously, we found that the activity-dependent epigenetic regulation of gene expression that this is associated with fear-related learning is modulated, in part, by nuclear lncRNAs ^12, 13^. We therefore wondered whether lncRNAs at the synapse are also a key feature of localised regulation of cellular processes underlying fear extinction. To address this, we used lncRNA capture-sequencing to map the expression of synapse-enriched lncRNAs in the ILPFC of adult male C57/bl6 mice, followed by single-molecule tracking in live cortical neurons and a CRISPR-inspired cell-type and synapse-specific and state-dependent RNA knockdown approach, which has revealed the importance of a novel variant of the lncRNA Gas5 that is critically involved in the trafficking of RNA granules at the synapse and in the formation of fear extinction memory.

## Results

### A significant number of lncRNAs are enriched at the synapse in the adult ILPFC

lncRNAs are predominantly expressed within nuclear subcompartments such as the nucleolus and in paraspeckles, where they have been relatively well characterised ^14-16^. To determine whether there are synapse specific lncRNAs that may be involved in fear extinction learning, we used synaptosome isolation followed by lncRNA capture sequencing on tissue derived from the adult mouse ILPFC (Figure 1a). To verify the purity of the synaptosome preparation, we used the dendritically localised scaffold protein PSD-95 as a marker (Supplemental Figure 1), and for targeted lncRNA enrichment, we employed a panel of 190,689 probes tiling across 28,228 known and predicted lncRNA transcript isoforms^17^. We identified 6970 lncRNA transcript isoforms that were expressed in both the nucleus ^13^ and synapse of ILPFC neurons (FPKM<0.5) (Figure 1b and 1d). Of these, 2583 were enriched in the synaptic compartment, including novel (62.5%, 1615) and annotated (GENCODE V25, 37.5%, 968) lncRNAs (Supplemental Table 1), with a significant proportion derived from intragenic (51.2%, 1324) and extragenic (28.7%, 741) regions (Figure 1c).

**Figure 1.**
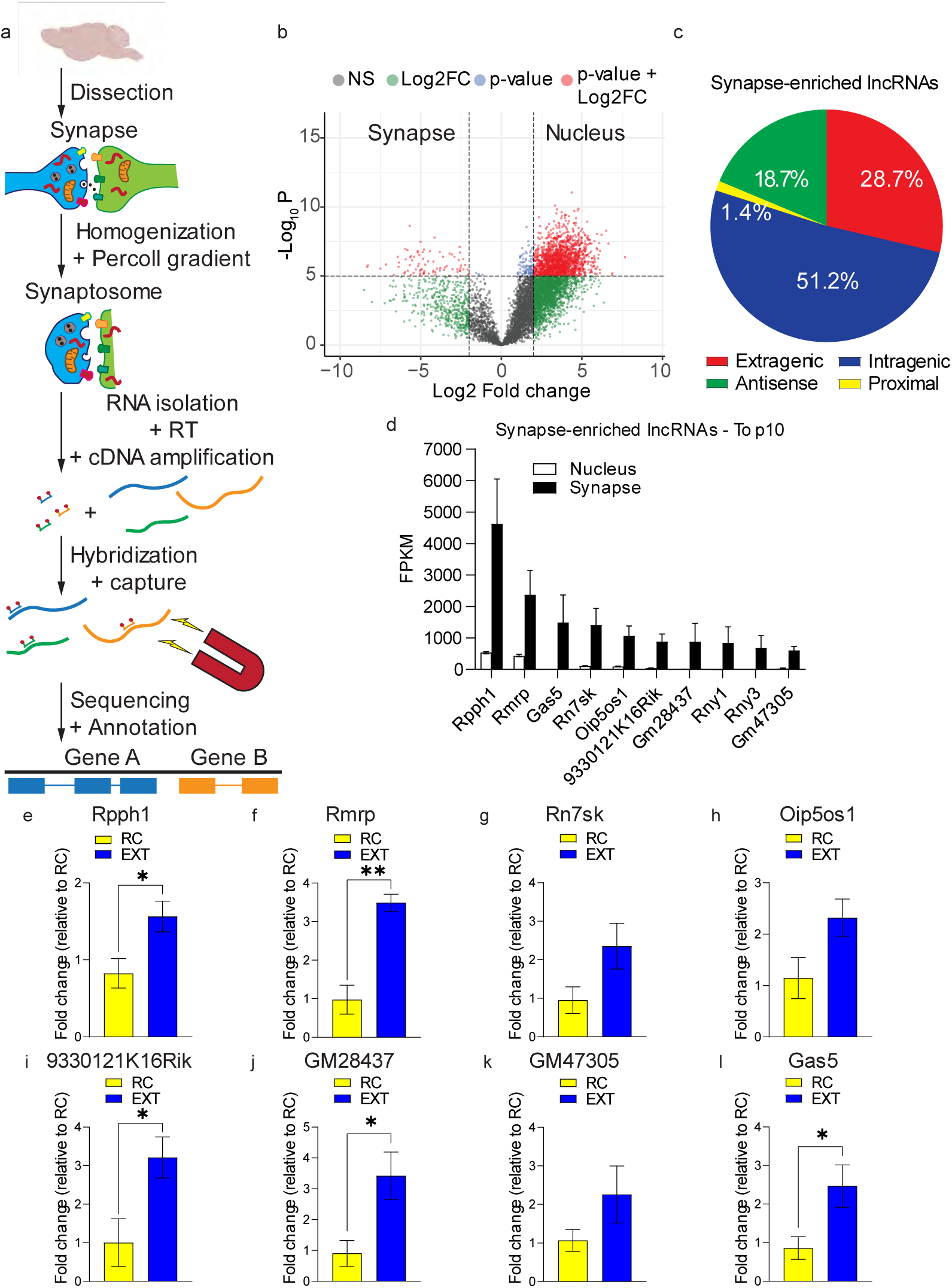
Targeted RNA capture-seq reveals a myriad of novel unannotated synapse-enriched lncRNAs. (A) Schematic overview of the synapse lncRNA capture-seq protocol. (B) Volcano plot showing all highly expressed (FPKM > 0.5) nuclear and synaptic lncRNAs in the ILPFC generated from the capture-seq data. The threshold for -log10 (*P*-value) set to 5 and the log2 (fold change) set to -2 and 2. Significant hits are highlighted in red. (C) Classification of captured nuclear and synaptic lncRNAs based on their genomic location with respect to protein coding genes, according to GENECODE annotation, version 25. (D) Bar plots showing the top 10 lncRNAs significantly expressed in the synapse. The reads are expressed as FPKM values. (E-L) RT-qPCR of 8 of the 10 candidates in (D) in the synapse in the ILPFC following fear extinction training. Bar plots are based on three or more biological replicates. Statistical significance was determined using two-tailed Student’s t-test. * p<0.05, ** p<0.01.

In contrast to our recent work on enhancer-derived lncRNAs (eRNAs) and memory, which revealed 434 eRNAs directly associated with fear extinction ^13^, we identified just 35 putative eRNAs at the synapse, suggesting that the majority of synapse-enriched lncRNAs are not likely to be involved in transcriptional regulation. Furthermore, the majority of synapse-enriched lncRNAs (76.9%, 1987) contained transposable elements, including both short interspersed nuclear elements (SINEs) and long interspersed nuclear elements (LINEs) (Supplemental Table 1), with a significant number found in antisense lncRNAs (18.7%, 483) (Figure 1c). These data suggest that the expression of transposable elements may be crucial for the functional activity of synapse-enriched lncRNAs. Indeed, recent evidence suggests that inverted antisense B2 SINEs or SINEUPs are important modulators of protein-coding gene translation ^18, 19^.

We next selected 8 of the top 10 synapse-enriched lncRNAs for validation by RT-qPCR (Figure 1e-l). Using an independent cohort of behaviourally-trained mice, a comparison between mice that had been fear conditioned followed by exposure to a novel context 24 hrs later with no further cue exposure (RC) and mice that had been fear conditioned followed by fear extinction training (EXT) revealed a fear extinction-learning induced increase in the expression of 5 lncRNAs, including, Rpph1, Rmrp, 93301121K16Rik, Gm28437 and Gas5 (Figure 1e-l). Among these, the stress responsive lncRNA *Gas5* has been implicated in the regulation of motivated behaviour ^20, 21^. Given that fear extinction is associated with changes in stress reactivity and involves prefrontal cortex-dependent behaviours, we decided to focus the remaining investigation on the functional relevance of *Gas5* in fear extinction.

### An alternatively spliced variant of Gas5 is enriched at the synapse

Upon closer analysis of the sequencing data, we found clear evidence for differential expression of Gas5 splice variants in the synapse compared to the nucleus ^13^ (Figure 2a). In contrast to intron-retained *Gas5* variants, which were confined to the nucleus, the *Gas5* variant ENSMUST00000162558.7 was preferentially localised at the synapse (Figure 2b and 2c, Supplemental Table 1). As a control, we also examined the localised expression of *Meg3* because it is activity-dependent, associated with synaptic plasticity, and previously shown to be enriched in the nucleus ^22^. As expected, the *Meg3* transcript was again preferentially localised to the nucleus (Figure 2c). Finally, the localised expression of the Gas5 variant was confirmed using BaseScope in primary cortical neurons, *in vitro* (Figure 2d). Our findings agree with the prevailing view that one important function of alternative splicing is to direct RNA localisation ^23, 24^, and extend this with evidence indicating that fear extinction learning leads to the accumulation of a specific Gas5 variant in the synaptic compartment.

**Figure 2.**
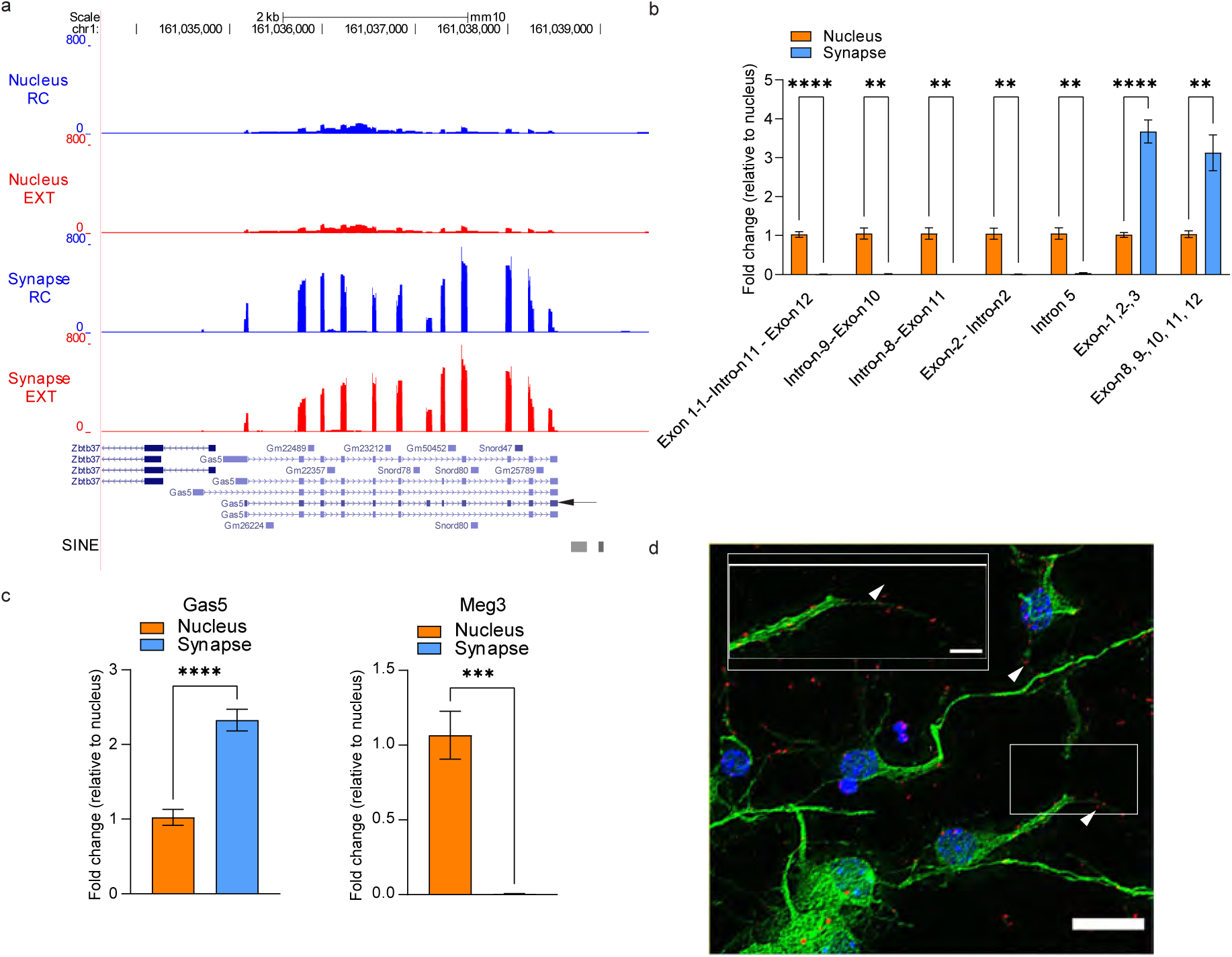
Gas5 is highly enriched in the synapse following fear extinction training. (A) Genomic tracks displaying lncRNA capture-seq data. Bars shown are reads distribution from nucleus and synapse RNAs derived from prefrontal cortex of retention control (RC) (blue) and fear-extinction trained (EXT) (red) mice. Transcripts are highlighted below the track. Grey bars indicate SINE elements across the region. Arrow indicates *Gas5* variant ENSMUST00000162558.7. (B) RT-qPCR of *Gas5* variants expression in the nuclear and synaptic fractions of the ILPFC. The amplified *Gas5* exonic and intronic regions are indicated. Bar plot is based on at least 4 biological replicates. Statistical significance was determined using two-tailed Student’s t-test. ** p<0.01, **** p<0.001. (C) RT-qPCR of *Gas5* variant ENSMUST00000162558.7 expression in the nucleus and synapse of the ILPFC region. Bar plot is based on at least 4 biological replicates. Statistical significance was determined using two-tailed Student’s t-tests. **** p<0.001. (D) Image displays synaptic *Gas5* variant in primary cortical neurons using BaseScope. Representative images from n > 8 fields of views. Arrowheads show synaptic localization. Scale bar, 20 µm. Red, *Gas5*; Blue, DAPI; green, MAP2 protein. The boxed region is enlarged in the inserts. Scale bar, 5 µm.

### The synapse-enriched *Gas5 variant* interacts with RNA binding proteins involved in translation and RNA localisation, as well as RNA granules

To begin to explore the Gas5-protein interaction network in the ILPFC and its relationship with behavioural experience, we performed RNA immunoprecipitation followed by mass spectrometry using a synthetically designed biotinylated Gas5 variant to pull down synapse-enriched lncRNA:protein complexes in ILPFC samples derived from RC and EXT mice. Overall, the Gas5 variant was observed to interact with 494 proteins at the synapse, with the majority (85%, 418) found in both RC and EXT groups. Within the RC group, we detected 67 (12%) unique proteins whereas 39 (8%) proteins were specific to EXT (Figure 3a). A String analysis of the global *Gas5*-protein network common to both groups revealed a significant cluster of RNA binding protein (RBP) interactions (Figure 3b). Next, a gene ontology (GO) analysis indicated a biological network associated with RNA processing (FDR 6.16e-55), which included molecular and cellular processes such as non-membrane organelles (light blue, FDR 9.05e-60), the spliceosome (dark blue, FDR 2.36e-20), the ribosome (red, FDR 6.19e-20) and an interaction with the cytoskeletal protein network (yellow, FDR 2.94e-13). These findings are consistent with the idea that RNA processing and the coordination of local translation is influenced by experience^25^. Importantly, several proteins associated with the non-membrane organelle cluster, including Caprin1, G3bp2, G3bp1, Pum1, Pum2, DDX6, Stau2, Atxn2 and Fxr1, have previously been associated with learning and memory ^26-29^. Together, these data suggest that local RNA metabolism may, in part by, be driven by the synapse-specific expression of lncRNA variants such as Gas5.

**Figure 3.**
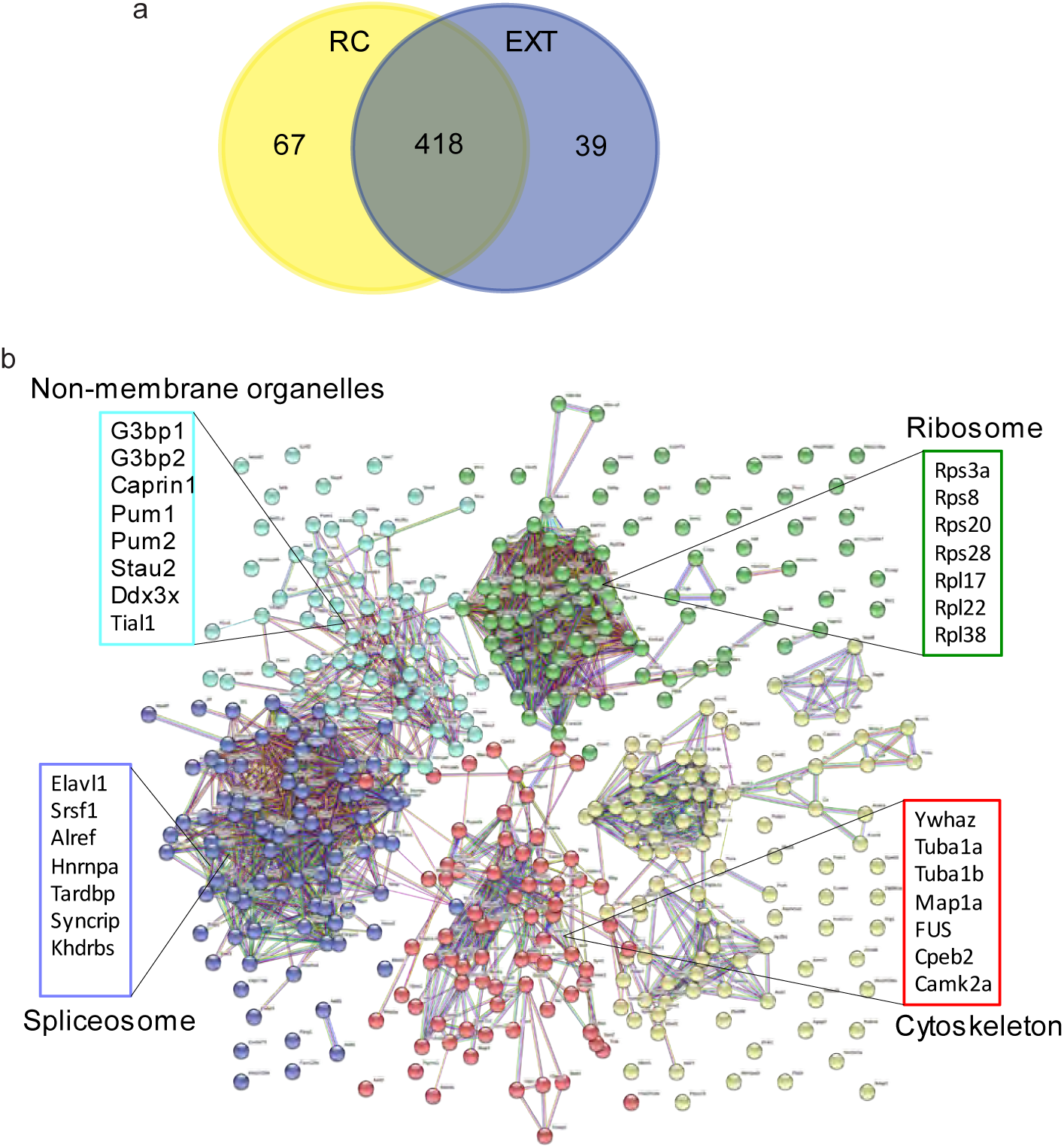
*Gas5* interacts with proteins involved in translation, and RNA localization, as well as RNA granules. (A) Venn diagram and enrichment of proteins that showed binding to *Gas5* in both RC and EXT groups are displayed. (B) STRING network analysis of the proteins that bound to *Gas5* in both RC and EXT groups are displayed. Only interactions with a STRING score ≥ 0.7 are shown. Evidence of interaction is represented by the distance between nodes, with more tightly packed nodes having a higher STRING score. In both groups, membraneless organelles, ribosome, cytoskeleton, and spliceosome clusters that contain tightly packed nodes are depicted. Samples were processed in biological triplicates.

We next selected Caprin1 and G3bp2 for further analysis because of their known role in the formation of RNA granules and in RNA trafficking ^28, 30^. To determine the structural module important for Caprin1 and G3bp2 binding, we generated a series of *Gas5* oligonucleotides with 50nt deletions tiled across a 502 nucleotide sequence spanning the splice sites for exons 1 through 12, and used these to immunoprecipitate Caprin1 or G3pb2 (Figure 4a). A decrease in Caprin1 binding was observed when the 3’ terminal end of *Gas5* (409-502 base pairs) was deleted, suggesting that this region of the Gas5 variant contains the module that is critical for the Gas5-Caprin1 interaction (Figure 4b and 4c). A modest decrease in Caprin1 binding was also observed when the region between nucleotides 204-308 was deleted. In contrast, a very significant reduction in G3bp2 binding occurred when the region between nucleotides 1-257 was deleted (Figure 4b and 4d). Whether the Gas5-Caprin1 or Gas5-G3bp2 interaction is regulated by RNA modification or dynamic changes in RNA structure remains to be determined. Nonetheless, these findings strongly suggest that there are specific regions of the Gas5 variant that are critical for a functional interaction between Gas5, Caprin1 and G3bp2.

**Figure 4.**
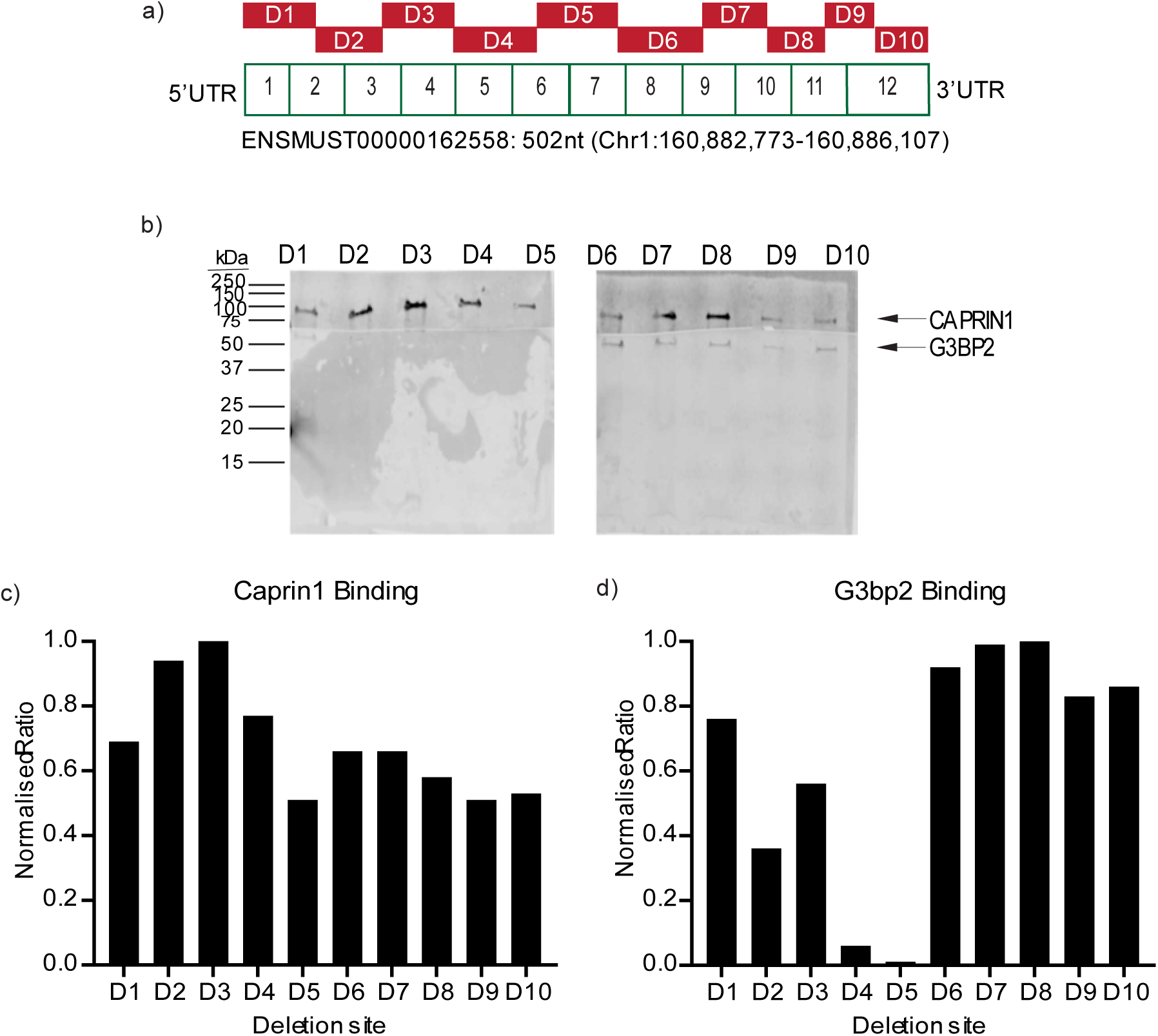
*Gas5* interacts with CAPRIN1 and G3BP2 using distinct structural modules. (A) Schematic showing deleted regions (D1 – D10) of *Gas5* RNA fragments used for *in-vitro* biotinylated RNA pull-down assay to identify the CAPRIN1- and G3BP2-binding regions on *Gas5*. (B) Blots displaying CAPRIN1 and G3BP2 proteins after incubating different fragments of *in-vitro* transcribed *Gas5* with ILPFC protein extracts. At least two independent experiments were performed. Normalized band intensity values of (C) CAPRIN1 and (D) G3BP2 are indicated.

### *Gas5* knockdown increases trafficking of RNA granules

Based on the finding that *Gas5* interacts with RBPs associated with RNA trafficking and RNA granules, including Caprin1 and G3bp2, we next investigated the functional relevance of this interaction by examining the effect of *Gas5* knockdown on RNA granule mobility using single-particle tracking photoactivation localization microscopy (sptPALM). We focused on G3bp2 because it forms a core complex with Caprin1 and is critically involved in RNA granule trafficking ^30^. To visualize and track RNA granules, G3bp2 was fused with the photoconvertible fluorescent protein mEos3.2, and then expressed under the control of the *Syn1* promoter, thereby allowing G3bp2-mEos3.2 specific expression in neurons. We then placed the whole cassette into a lentiviral backbone, packaged the virus and transfected primary cortical neurons. Further, given that the expression of the *Gas5* variant is increased at the synapse following fear extinction training, we investigated the functional consequence of Gas5 variant knockdown at the synapse.

To decrease the expression of the this variant selectively at the synapse, we used a CRISPR-Cas-inspired RNA targeting system (CIRTS), which is a guide RNA (gRNA)-dependent technology designed to deliver protein cargoes to target RNAs, as its small size facilitates viral packaging and protein delivery to the brain ^31^. To target the PIN nuclease effector for RNA degradation at the synapse, we appended the full length *Calm3* intron as a dendritic localisation signal ^32^ and expressed the CIRTS cassette under the control of a neuron-specific *Syn1* promoter on a lentiviral backbone (Figure 5a). For visualization, GFP was fused upstream of the CIRTS cassette together with a 2A self-cleaving peptide signal. To test the functionality of this system, we first designed two gRNAs to specifically target the synaptic *Gas5* variant ENSMUST00000162558.7 and examined the effect of each guide in primary cortical neurons. One of these guides degraded *Gas5* variant ENSMUST00000162558.7 by more than 50%, and was therefore chosen for all subsequent knockdown experiments (Figure 5c). We also determined whether the CIRTS-Gas5 construct localised to dendrites in a KCl-induced chase experiment in primary cortical neurons. After a 10-minute pulse and chase for 5 minutes, an increase in localised CIRTS-Gas5 expression was observed in the dendritic spines of primary cortical neurons, with the accumulation increasing 60 minutes after stimulus (Figure 5b), thereby indicating that our construct localises to the synapse in an activity-dependent manner.

**Figure 5.**
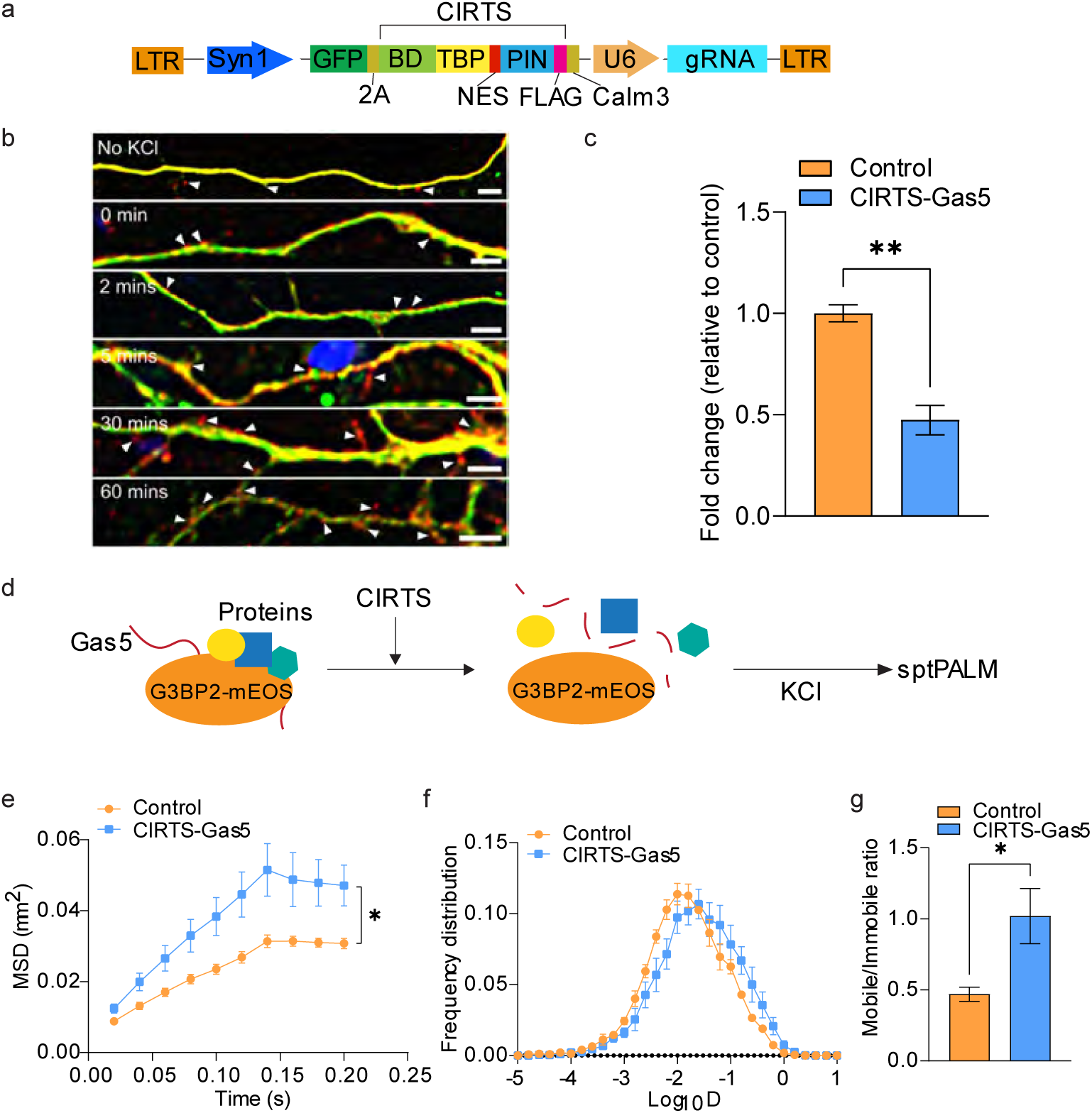
*Gas5* knockdown impairs the mobility of RNA granules. (A) Schematic representation of the viral CIRTS construct used for *Gas5* knockdown in primary cortical neurons. Syn1 and U6 promoters drive the expression of the CIRTS and guide RNAs, respectively. GFP is inserted upstream of CIRTS for visualization. NES, nuclear export signal; Calm3, Calm3 dendritic localization signal; 2A, self-cleaving peptide signal. (B) Immunofluorescence detection of CIRTS in dendrites in primary cortical neurons. Time after 10 min of KCl induction is indicated. Arrow heads show CIRTS puncta (Red)in the dendritic spines. Representative images from n > 8 field of views. Scale bar = 5 µm. Green, MAP2; Red, CIRTS. (C) qRT-PCR validation performed on primary cortical neurons transduced with either control or CIRTS-*Gas5* virus. N >2 biological replicates. Statistical significance was determined using two-tailed Student’s t-test. * p<0.05. (D) Schematic diagram of G3BP2-mEos3.2 sptPALM analysis. G3BP2-mEos3.2 was excited by 561 nm laser and photoconverted by 405 nm laser in the presence of dendritic-localised CIRTS control or CIRTS-*Gas5* and KCl. Proteins and *Gas5* RNA that bind to G3BP2 are indicated. (E-G) Change in mean squared displacement (MSD) (µm^2^), relative frequency distribution of diffusion coefficients and ratio of mobile to immobile fractions of G3BP2-mEos3.2 elicited on CIRTS-*Gas5* or control knockdown under KCl. At least 3 neurons were analysed per group. An average of 1678 and 2054 trajectories per neuron in the control and CIRTS-*Gas5* samples respectively, were analysed. Statistical tests were performed using the Student’s t-test (two-tailed distribution, unpaired). * p<0.05, Means ± s.e.m. are plotted.

We next expressed G3bp2-mEos3.2 in the presence of either a CIRTS scrambled control or CIRTS-*Gas5* in primary cortical neurons and subjected them to sptPALM analysis (Figure 5d). To quantify G3bp2 mobility, we analysed the mean square displacement (MSD) of the trajectories of individual G3bp2-mEos3.2 molecules obtained by sptPALM, to examine changes in mobility in the presence of the scrambled control or CIRTS-calm3-*Gas5*. A comparison of the MSD between knockdown and control conditions under low KCl stimulation conditions revealed a significant increase in mobility following *Gas5* knockdown as evidenced by a less constrained curve (Figure 5e). Analysis of the diffusion coefficient distribution of G3bp2-mEos3.2 molecules revealed a shift in the mobile and immobile fractions (Figure 5f). This was apparent when we compared the mobile:immobile ratio (0.47 and 1.02 for control and *Gas5* knockdown, respectively) (Figure 5g). Our data suggest that *Gas5* controls the nanoscale organisation of G3bp2, which then coordinates the activity RNA granules at the synapse.

### Targeted knockdown of the synapse-enriched *Gas5* variant in the ILPFC impairs fear extinction memory

To assess the functional role of *Gas5* activity at the synapse in fear extinction, we infused the synapse-targeted CIRTS-Gas5 construct into the ILPFC prior to training (Figure 6a and 6d). The knockdown efficiency *in vivo* was first validated by RT-qPCR (Figure 6b and 6c), which showed a selective reduction in Gas5 in the synaptic compartment with little effect on nuclear Gas5 expression. There was no effect of Gas5 knockdown on within-session fear extinction training (Figure 6e) nor was there any effect on the ability to express fear in the absence of fear extinction training when tested 24 hours after training (Figure 6f, white vs black bars). In contrast, Gas5 knockdown led to a significant impairment in fear extinction memory (Figure 6f, yellow vs blue bars). To determine the effect of *Gas5* knockdown on the stability of the original fear memory, the mice were re-exposed to the original training context A 24 hours later. There was no significant difference between RC Control and EXT CIRTS-*Gas5* animals in fear expression (Supplemental Figure 2), suggesting that *Gas5* knockdown did not alter fear memory per se. Collectively, these data indicate that synapse-directed *Gas5* knockdown in the ILPFC selectively impairs fear extinction memory without interfering with the original fear memory trace.

**Figure 6.**
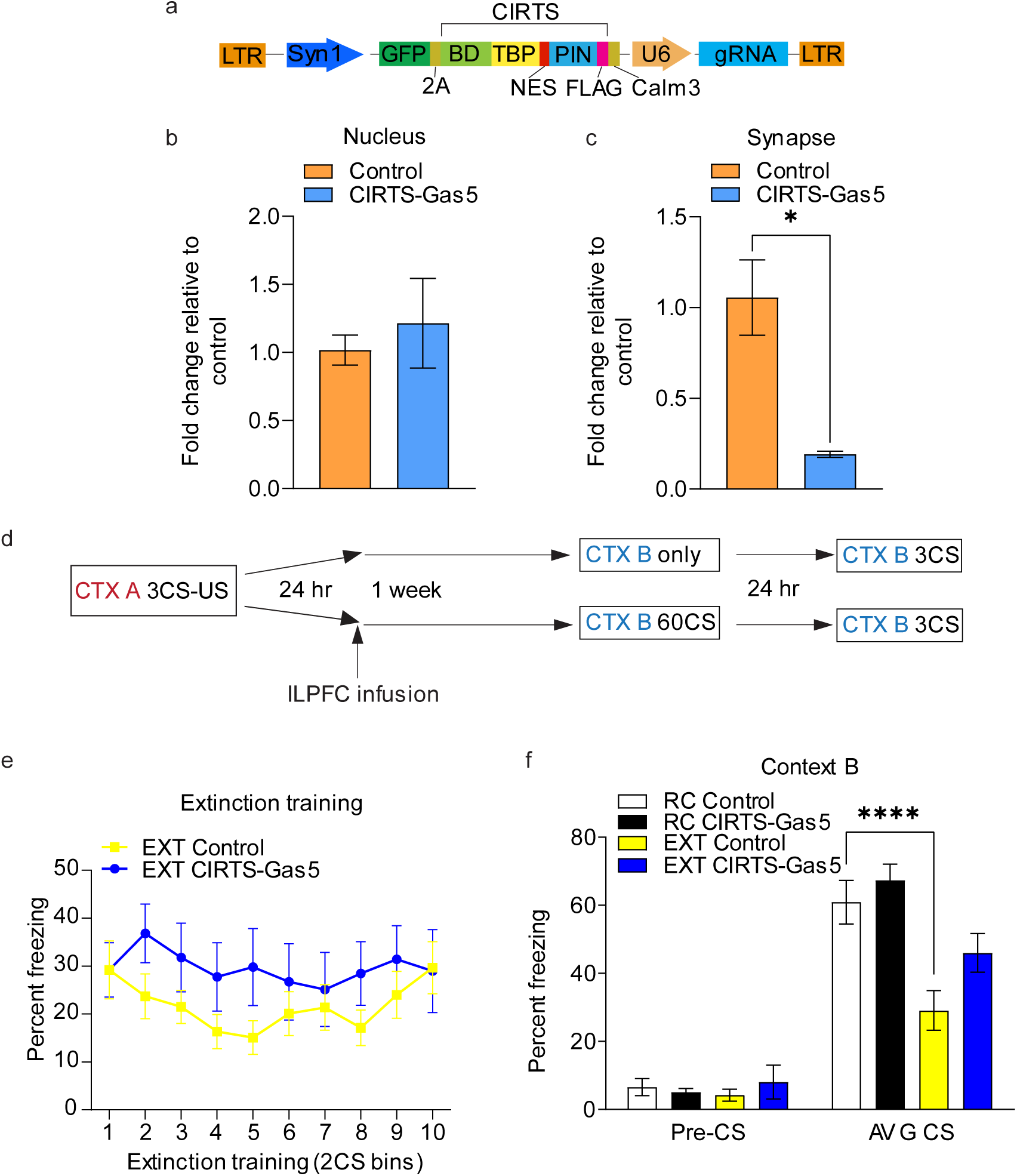
*Gas5* knockdown impairs fear extinction memory. (A) Schematic representation of the viral CIRTS construct used for Gas5 knockdown in the ILPFC in mice. Syn1 and U6 promoters drive the expression of the CIRTS and guide RNAs, respectively. GFP is inserted upstream of CIRTS for visualization. NES, nuclear export signal; DLS, Calm3 dendritic localization signal; 2A, self-cleaving peptide signal. qRT-PCR verification of Gas5 knockdown in the nucleus (B) and at the synapse (C). Box plot is shown based on at least two biological replicates. Statistical significance was determined using the two-tailed Student’s t-test. * p<0.05. (D) Schematic of the behavioural protocol used to test the effect of Gas5 knockdown in the ILPFC on fear extinction memory. CTX, context; CS, conditioned stimulus; US, unconditioned stimulus. (E) Graph showing within-session performance during the first 10 conditioned stimulus exposures during fear extinction training (n > 5 biologically independent animals per group, two-way ANOVA with Bonferroni’s post hoc test, all p>0.9999) (F) Bar plot showing animals treated with control virus, fear conditioned and exposed to a novel context (RC control); control animals fear conditioned followed by extinction (EXT control); Gas5 knockdown animals fear conditioned and exposed to a novel context (RC CIRTS-Gas5); and Gas5 knockdown animals fear conditioned followed by extinction (EXT CIRTS-Gas5); n > 5 biologically independent animals per group, repeated two-way ANOVA with Dunnett’s posthoc tests. **** p <0.001

## Discussion

In this study, we have revealed a significant population of lncRNAs at the synapse and discovered a localised isoform of *Gas5* that is functionally active and required for fear extinction memory. This *Gas5* variant interacts with proteins involved in translation, RNA trafficking, RNA metabolism and the formation of RNA granules at the synapse, including Caprin1 and G3bp2. Together, these observations suggest a novel mechanism by which lncRNA activity at the synapse can influence the behaviour of RNA condensates through the binding of granule proteins. Gas5 may serve as a guide to coordinate the trafficking of RNA granules, and to organize the learning-induced activity of key proteins involved in local protein translation and memory formation (Figure 7).

**Figure 7.**
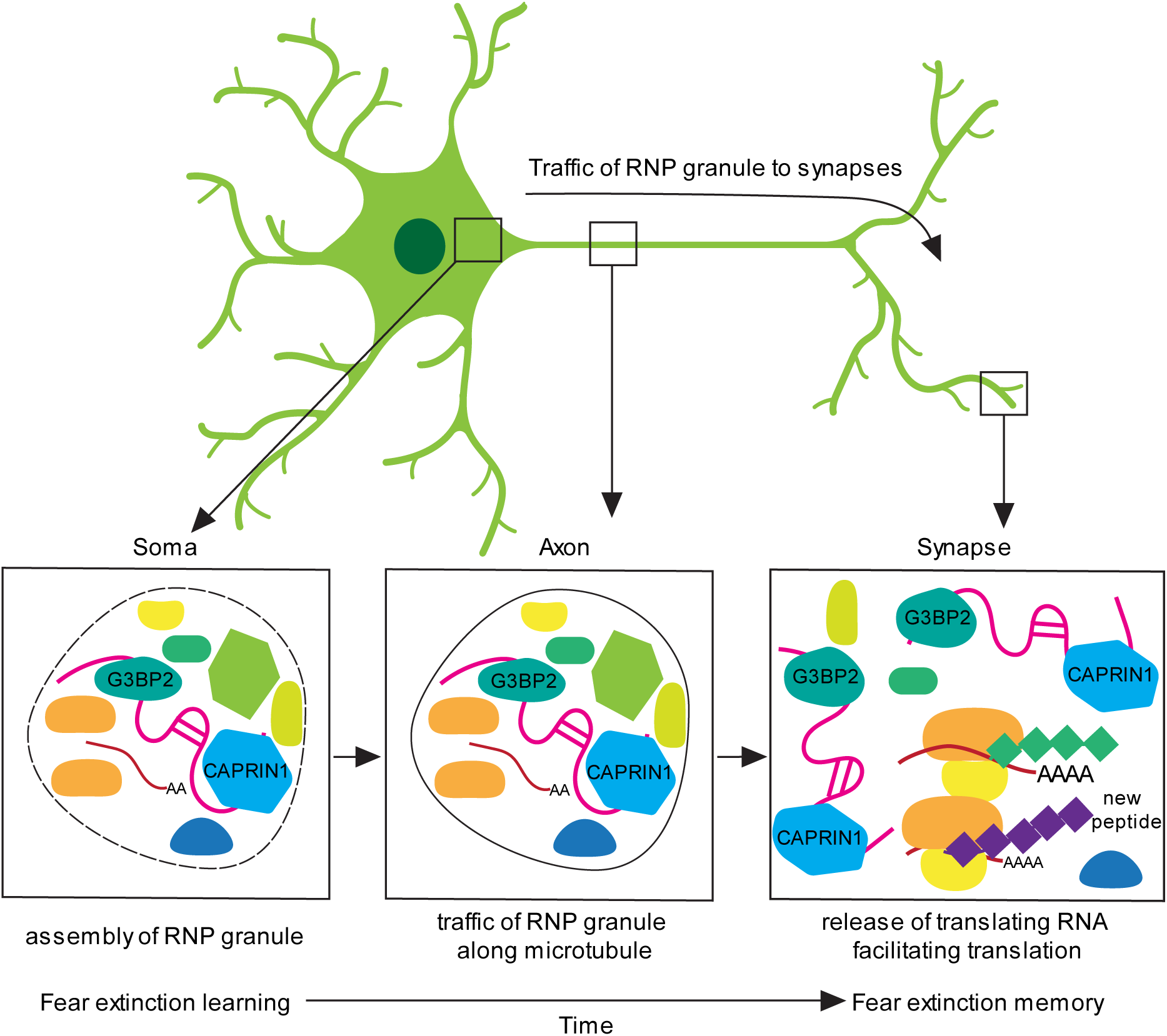
A model for the mechanism by which *Gas5* coordinates RNA granules at the synapse during fear extinction learning. Extinction learning leads to the assembly and transport of CAPRIN1-G3BP2-*Gas5* RNA granules to the synapse and the accumulation of *Gas5* variant in the synaptic compartment sequesters CAPRIN1 and G3BP2 from binding to translating RNAs facilitating local protein synthesis.

The *Gas5* lncRNA undergoes a significant degree of alternative splicing, yielding 25 isoforms. Our findings indicate that the synaptic *Gas5* variant in the ILPFC is a mature isoform that is functionally involved in fear extinction memory. It is interesting to note that a different splice variant of *Gas5* has been detected in the nucleus accumbens and plays a crucial role in motivated behaviour^21^, suggesting a role for alternative splicing in brain-region specific expression of the Gas5 lncRNA. Most Gas5 exons are flanked by small nucleolar RNA (snoRNAs) found within their introns, a unique feature found in sno-lncRNAs^33^. We found that the intron-retained transcripts are nucleus-specific, suggesting that these transcripts haven’t had their introns removed at this timepoint and are thus processing intermediates. As most Gas5 transcripts have at least one snoRNA, they may also function as guides for RNA modification in the nucleus. Indeed, intron-retained, poorly spliced lncRNAs are more commonly found in the nucleus^34^. In addition to a multifunctional role for Gas5 as a sno-lncRNA, it also remains possible that the *Gas5* lncRNA is capable of both coding and coding-independent functions, similar to that described for coding and noncoding RNAs (cncRNAs)^35, 36^. In support of this idea, the human ortholog of *Gas5* is just one of many lncRNAs that contain short open reading frames (sORFs) that may yield microproteins that are localised to different subcellular compartments and involved in diverse cellular functions^37^. Future studies will investigate whether there are sORFs derived from the Gas5 locus that are translated and functionally active in the adult brain.

Previous work has shown that lncRNAs can associate with ribonucleoproteins (RNPs) and other RNAs to form RNA condensates. For example, neuroLNC can regulate TDP-43 granule localization to the synapse ^38, 39^. In addition, early studies also suggested that Caprin1-containing granules are localised to dendrites and important for long-term memory formation ^28^. The formation of RNA granules is therefore dependent on the properties of both RNA and protein and the surrounding microenvironment^40, 41^. We found that the “RNP complex” was the most abundant protein category in the RNA-protein network with proteins involved in RNA granule formation being the most prominent. Our findings demonstrate that the synapse-enriched *Gas5* variant binds to the RNA granule proteins, Caprin1 and G3bp2. These proteins are known to form a stable complex in response to stress ^30^, which serves to regulate condensate localization and dynamics. Caprin1-containing granules are also important for AMPA receptor loading and dendritic RNA localization in postsynaptic neurons ^28^. We found that Gas5 knockdown affects the mobility and trafficking of G3bp2-containing RNA granules, suggesting that Gas5 acts as a scaffold to coordinate the mobility of RNA granules in an activity-dependent manner. It is therefore plausible that the destabilisation of these “memory” granules might represent a critical factor associated with impairments in the formation of long-term extinction memory.

Finally, many of the identified synapse-enriched lncRNAs were unannotated or labeled as pseudogenes with no function described. The term “pseudogene” was first used to categorize non-functional duplicated genes ^42^. However, recent studies have shown that duplicated pseudogenes may be lncRNAs ^43^. Indeed, some pseudogene-derived lncRNAs are induced in an activity-dependent manner and are required to maintain synaptic plasticity ^44^. Interestingly, we found that many pseudogenic lncRNAs contain LINE and SINE elements, suggesting that these elements may be associated with their function. Recent studies have also shown that SINE- and LINE-containing lncRNAs are responsible for translation and RNA localization and are often isoform-specific ^18, 19^. Given that lncRNAs have been shown to contain various structural modules that can also serve as a decoy for microRNAs ^45^, as guides for axon regeneration ^46^ or as scaffolds for ribonucleoprotein localization ^47, 48^, and the fact that dual functioning lncRNAs are increasingly becoming the norm rather than the exception ^12, 49, 50^, future studies should aim to dissect the structure-function relationship of each module, determine its conservation, and reveal its role in neuronal function.

The work presented here leverages the synapse-enriched lncRNA capture-seq approach. However, because of input requirements and the amount of RNA that can be derived from the synaptic compartment, we were unable to achieve cell-type resolution. Given that the current lncRNA annotations are incomplete, it is also possible that other lncRNAs beyond those reported here are present at the synapse during fear extinction learning. Increasing the sample input and further refinement of the lncRNA set targeted by the probes are needed to capture the full spectra of synapse-enriched lncRNAs in the ILPFC. Although we have functionally characterised a lncRNA that is required for the mobility of one type of granule in the dendrite, this mechanism may not hold true for other lncRNAs or other granules. Further work is needed to explore the functional and mechanistic roles of individual synaptic lncRNAs and their interacting partners in different state-dependent conditions and in other learning paradigms.

In conclusion, we have discovered a synapse-enriched lncRNA isoform that is associated with the formation of fear extinction memory. We have discovered that through the regulation of RNA granule trafficking at the synapse the localised expression of a variant of *Gas5* is required for fear extinction memory. Our data support the idea that the coordinated activity of RNA granules is required for long-term memory formation and place lncRNA-mediated regulation of RNA condensates as an important factor underlying the formation of extinction memory.

## Supporting information

Supplemental Tabl3

## Acknowledgments

We thank Dr. Alun Jones from the Mass Spectrometry Facility in the Institute of Molecular Bioscience at the University of Queensland for help with the proteomics experiments and Ms. Rowan Tweedale in the QBI for help in manuscript editing. The authors gratefully acknowledge the QBI Advanced Microscopy Facility for their support and assistance in this work. We thank JB Sibarita (University of Bordeaux, France) for providing the PalmTracer software. The authors acknowledge grant support from the Brain and Behavioural Research Foundation (NARSAD Independent Investigator Award, T.W.B.), NIH R01MH109588 (TWB and RCS), NHMRC Ideas Grant (GNT2003414), NSFC 82001421 (X.L.), the National Institute of General Medical Sciences of the National Institutes of Health NIH (R35 GM119840 to B.C.D.) and DFG (SFB870, SPP1935, M.A.K), as well as the Australian Research Council Discovery Grant DP190100674 (F.A.M) and a National Health and Medical Research Council (NHMRC) NHMRC Senior Research Fellowship (GNT1155794) to F.A.M.). E.L.Z, L.J.L. and S.U.M. are supported by a Westpac Future Scholarship and the University of Queensland.

## Author Contributions

W. L. conceived this project, together with T.W.B. and R.C.S., and led the development and optimization of the protocol, performed experiments, analysed and interpreted data, generated figures, and wrote and edited the paper. M.C. shared the lncRNA probe set. H.G., L.J.L., S.U.M., A.P., X.L. W.W. and E.L.Z. assisted with tissue collection, surgery and behavioural experiments. P.R.M. performed the behavioural experiments and analysis. B.A. performed sptPALM experiments and B.A. R.S.G analysed the data. F.A.M. supervised the sptPALM analysis. Q.Z. performed bioinformatics analysis. All authors discussed results and revised the manuscript.

## Methods

### Animals

Male C57BL/6 mice (10–14 weeks old) were housed two per cage, maintained on a 12 h light/dark time schedule and allowed free access to food and water. All testing took place during the light phase in red-light-illuminated testing rooms following protocols approved by the Institutional Animal Care and Use Committee of the University of California, Irvine and by the Animal Ethics Committee of The University of Queensland. Animal experiments were carried out in accordance with the Australian Code for the Care and Use of Animals for Scientific Purposes (8th edition, revised 2013).

### Plasmid Construction

The pFsy(1.1)GW lentiviral expression vector (addgene #27232), containing the synapsin 1 promoter, was used to make the Syn1-Cirts-Calm3 construct. The CIRTs cassette was PCR amplified and cloned into the AgeI and XbaI site. A NheI site was generated upstream of XbaI using PCR primer. The calm3 dendritic localization sequence was PCR amplified from psiCheck2-Calm3 and cloned into the NheI and XbaI site. The U6-gRNA was PCR amplified from the CIRTS pin nuclease construct and inserted in the XbaI site. GFP was PCR amplified and cloned into the AgeI site along with a 2A peptide signal. G3BP2-mEos3.2 were constructed by fusing mEos3.2 to the C-terminus of G3BP2. The fragment was PCR amplified and inserted into the AgeI and XbaI sites of the pFsy(1.1)GW lentiviral vector.

### Tissue Culture

Cortical tissue was isolated from embryonic day 16-17 C57BL/6 embryos. Primary cortical neurons were isolated by removing the skull and meninges with fine-tipped tweezers. Cells were then mixed with Neurobasal medium (Gibco 21103) containing 5% fetal bovine serum (FBS), B27 supplement (Gibco 17504-044), GlutaMAX (Gibco 35050) and 1% penicillin-streptomycin (Gibco 15140) and made homogenous with gentle pipetting. Cells were then passed through a 40 µm cell strainer (BD Falcon 352340) and plated onto culture dishes coated with poly-L-Ornithine (Sigma P2533). HEK293T cells were maintained in a medium containing Dulbecco’s modified eagle medium (DMEM) and high glucose (Gibco 11965) with 5% FBS and 1% penicillin-streptomycin.

### Synaptosome Preparation

Preparation of synaptosomes was carried out as previously described ^51^. Briefly, tissues were homogenized in homogenizing buffer with 10-14 strokes using a Teflon-glass tissue grinder. The homogenate was centrifuged at 1000 x g for 10 mins at 4°C. The supernatant was directly applied onto a discontinuous Percoll gradient ranging from 0% up to 23% Percoll and centrifuged at 31,000 x g for 5 mins to isolate the synaptosome fraction.

### RNA Extraction

Cultured cells, synaptosomes and tissues were homogenized using a dounce tissue grinder in NucleoZOL (Macherey-Nagel 740404) supplemented with 5 mM of DTT (Thermo Scientific R0862) and 0.1 mM of RNAseOUT (Invitrogen 10777019). RNA was extracted according to the manufacturer’s instructions. The concentration of RNA was measured using a nanophotometer (IMPLEN) or Qubit fluorometer (Thermo Fisher Scientific Q33216).

### RT–qPCR

1 µg of RNA was used for cDNA synthesis using the QuantiTect Reverse Transcription Kit according to the manufacturers’ protocol (Qiagen 205313). Quantitative PCR was carried out on a RotorGeneQ (Qiagen) real-time PCR cycler with SensiFAST SYBR master mix (Bioline BIO-98050) using primers for target genes and 18S rRNA or phosphoglycerate kinase (PGK) as an internal control. All transcript levels were normalized to 18S rRNA or PGK using the ΔΔCT method and each PCR reaction was run in duplicate for each sample and repeated at least twice.

### Western Blot

Homogenized issue and synaptosomes were fractionated in NP40 cell lysis buffer (Thermo Fisher Scientific FNN0021). Briefly, samples were incubated on ice in lysis buffer for 30 mins and then centrifuged at 14,000 rpm for 10 mins at 4°C. The supernatant was transferred to a new tube and protein concentration was measured with Bradford assay (Sigma B6916) on a nanophotometer (IMPLEN). Protein was diluted in Laemmli sample buffer with 5% 2-Mercaptoethanol (Sigma-Aldrich M6250) and denatured for 5 mins at 95°C. Gels were run and proteins transferred onto PVDF membrane (BioRad 1620264). The membrane was blocked with Odyssey Blocking Buffer (Li-Cor LCR-927-40003) for 1 hr at room temperature and incubated with primary antibody overnight at 4°C. Primary antibodies used were anti-Caprin1 (Proteintech 151121AP), G3bp2 (Abcam ab86135), PSD95 (Abcam ab13552), B-actin (Cell signalling 3700). The membrane was washed with phosphate buffered saline containing 0.2% Triton X-100 (PBST) (3x), incubated for 1 hr with IRDye 800CW secondary antibody (Li-COR) at 1:15,000 in PBST, and washed in PBST for 10 min (3x). Optical density readings of the blots were made using the LI-COR analysis system.

### Lentiviral Production

Plasmid was co-transfected with pMD2.G (Addgene #12259), pRSV-Rev (Addgene #12253) and pMDLg/pRRE (Addgene 12251) into HEK293T cells at approximately 80% confluence using Lipofectamine 3000 transfection reagent (Thermo Fisher Scientific L3000150). 4 hrs later, sodium butyrate (Sigma 303410) was added to stimulate viral production. After 2 days’ incubation at 37 °C and 5% CO_2_, the virus was collected by ultracentrifugation. The titer was measured using a Lenti-X qRT-PCR titration kit (Clontech 631235).

### Lentiviral Infusion and Behavioural Analysis

Lentiviral delivery and behaviour analysis were performed as reported in Marshall et. al. (2020). Briefly, a double cannula (PlasticsOne) was implanted into the ILPFC. Animals were then separated into single housing and given at least one week to recover prior to behavioural training. Mouse were first fear-conditioned in context A, followed by 2 lentiviral infusions 24 hrs post-fear conditioning, and after a one-week incubation, mice were either extinction-trained or just exposed to a novel context (context B). Movement was captured by cameras within the boxes and were processed automatically with FreezeFrame program (Actimetrics). The fear-conditioned protocol consisted of a 120 sec pre-fear conditioning incubation followed by three pairings of 120 sec, co-terminating with a 1 sec 0.7 mA foot shock (unconditioned stimulus, US). For extinction training, mice were exposed in context B in which they habituated to chamber for 2 min and extinction training (EXT) comprised of 60 non-reinforced 120 sec CS presentations (5-sec intervals). For the retention control (RC) animals, context exposure was performed as in the EXT group but without presentation of the tones. Freezing was assessed during three 120 sec conditioned stimulus (CS) presentations (120 sec interval). Memory was inferred by the percentage of freezing during the tests.

### Immunohistochemistry

Animals were perfused with 4% paraformaldehyde and brains were collected in 30% sucrose prior to slicing. Sectioning at 40 µm was performed using a vibratome. Sections were incubated 1 hr in blocking buffer and incubated with primary antibody at 4 °C overnight. Slices were washed with PBST (3x), after which secondary antibodies were added. Slices were incubated at room temperature for 1 hr, washed 3 times with PBST and mounted on Superfrost Plus microscope slides (Thermo Fisher Scientific 22-037-246) with ProLong Gold Antifade Mountant (Thermo Fisher Scientific P36982) with 4′,6-diamidino-2-phenylindole (DAPI) (Thermo Fisher Scientific D1306). Sections were imaged with a slide scanning confocal microscope.

### Immunofluorescence

Primary cortical neurons were fixed in 10% neutral buffered formalin solution (Sigma-Aldrich HT5011) at room temperature for 30 mins, after which they were washed in PBS (3x) and incubated with 4% goat serum in PBST at room temperature for 1 hr. Cells were then incubated with primary antibody at 4 °C overnight, washed with PBST (3x), and incubated with secondary antibody at room temperature for 1 hr. Finally, cells were washed in PBS (3x), stained with DAPI and mounted on Superfrost Plus microscope slides (Thermo Fisher Scientific 22-037-246) with ProLong Gold Antifade Mountant (Thermo Fisher Scientific P36982). Primary antibodies used were anti-Map2 (Abcam ab5392) and anti-GFP (Abcam ab6556). Secondary antibodies were anti-chicken Alexa Fluor 488 (Thermo Fisher Scientific A11039) and anti-rabbit Alexa Fluor 546 (Thermo Fisher Scientific A11035).

### RNAscope

Primary neurons were processed based on the manufacturer’s instructions for the Basescope RED assay (Advanced Cell Diagnostics ADV323910) with few modifications and combined with an immunofluorescence protocol. First, samples were subjected to protease treatment, probe hybridization (RNAscope probe BA-Mm-Gas5-tv224-E8E9 ACD ADV833211), amplification and signal development. Blocking buffer containing 4% goat serum in PBST was added and incubated at room temperature for 1 hr. Samples were then stained with primary and secondary antibodies (as described in the immunofluorescence protocol above). Amplified signal was detected using the cyanine 5 channel.

### Image Acquisition and Analysis

Tissue and primary neurons were imaged on an Axio Imager Z1 upright fluorescence microscope (Carl Zeiss) fitted with an Axiocam MRm camera (Carl Zeiss) and a 40X/0.75 NA Plan-Apochromat objective. Image acquisition was performed using ZEN software (Carl Zeiss). For tissue section, a 2.5X/0.075NA Plan-Apochromat objective was used. Images were analysed using ImageJ and figures were constructed using the FigureJ plugin ^52^.

### lncRNA Capture Sequencing

Mouse tissue and synaptosomes were collected and total RNA was isolated as described above. cDNA libraries were generated using NEBNext Ultra II RNA Library Prep Kit for Illumina (NEB E7645) according to the manufacturer’s protocol. A custom-designed probe panel (ROCHE) targeting 28,228 mouse lncRNAs ^17^ was used to capture amplified cDNA. The capture procedure was performed using the SeqCap EZ Hybridization and Wash Kit (ROCHE 05634261001) and SeqCap EZ Accessory kit (ROCHE 07145594001) according to the manufacturer’s instructions. Captured libraries were sequenced on an Illumina HiSeq 4000 platform with 150-bp paired-end reads (GENEWIZ).

### Sequencing Data Analysis

Cutadapt (v1.17, https://cutadapt.readthedocs.io/en/stable/) was used to trim low-quality nucleotides (Phred quality lower than 20) and Illumina adaptor sequences at the 3’ end of each read for both lncRNA capture sequencing data. Processed reads were aligned to the mouse reference genome (mm10) using HISAT2 (v2.1.0) (Kim et al, 2015). SAMtools (version 1.8) ^53^ was then used to convert “SAM” files to “BAM” files, remove duplicate reads, and sort and index the “BAM” files. To avoid the artefact signals potentially introduced by misalignments, we only kept properly paired-end aligned reads with a mapping quality of at least 20 for downstream analyses. Three rounds of StringTie (v2.1.4) ^54^ was applied to i) perform reference-guided transcriptome assembly by supplying the GENCODE annotation file (V25) with the “-G” option for each sample, ii) generate a non-redundant set of transcripts using the StringTie merge mode, and iii) quantifying the transcript-level expression for each sample, with the option of “-e -G merged.gtf”. Known protein-coding transcripts (with the GENCODE biotype as “protein_coding”) were removed from the StringTie results. Ballgown (v2.22.0) ^55^ was used to conduct transcript-level differential expression analysis.

### RNA Pull Down Assay

*Gas5* variants were amplified using the T7 promoter sequence on the 5’ end of the forward primers. The PCR products were gel extracted using the Gel DNA Recovery Kit (Zymo Research D4008) and *in-vitro* transcribed using the HiScribe T7 Quick High Yield RNA Synthesis Kit (NEB E2050) according to the manufacturer’s instructions. The transcribed RNA was purified using the RNA Clean and Concentrator Kits (Zymo Research R1014) and pull down was performed using the Pierce Magnetic RNA-Protein Pull-Down Kit (Thermo Fisher 20164) according to the manufacturer’s instructions.

### HPLC/MS MS/MS, Mass Spectrometry and Protein Identification

Magnetic affinity beads were covered with 40 µl of 40 ng/µl sequence grade trypsin in 50mM ammonium bicarbonate pH8 buffer (Promega V5111). The beads were placed in an incubator at 37C overnight. The trypsin solution was removed from each sample and placed in a clean Eppendorf tube. 200 µl of 5% formic acid/acetonitrile (3:1 (vol/vol) was added to each tube and incubated for 30 mins at room temperature in a shaker. The supernatant was placed into the pre-cleaned Eppendorf tubes, together with the trypsin solution for each sample and dried down in a vacuum centrifuge.

For HPLC/MS MS/MS analysis, 15 µl of 1.0% (vol/vol) TFA in water was added to the tube, which was vortexed and/or incubated for 2 min in the sonication bath and finally, transferred to an autosampler vial for analysis. Tryptic peptide extracts were analyzed by microflow HPLC/MS MS/MS on an Eksigent, Ekspert nano LC400 uHPLC (SCIEX, Canada) coupled to a Triple TOF 6600 mass spectrometer (SCIEX) equipped with a micro Duo IonSpray, ion source. 5 µl of each extract was injected onto a 5mm x 300 µm, C18 3 µm trap column (SGE,) for 6 mins at 10 µl/min. The trapped tryptic peptide extracts were then washed onto the analytical 300 µm x 150 mm Zorbax 300SB-C18 3.5 µm column (Agilent Technologies) at 3 µl/min and a column temperature of 45°C. Linear gradients of 2-25% solvent B over 60 min at 3 µl/min flow rate, followed by a steeper gradient from 25% to 35% solvent B in 13 min, then 35% to 80% solvent B in 2 mins, were used for peptide elution. The gradient was then returned to 2% solvent B for equilibration prior to the next sample injection. Solvent A consisted of 0.1% formic acid in water and solvent B contained 0.1% formic acid in acetonitrile. The micro ionspray voltage was set to 5500V, declustering potential (DP) 80V, curtain gas flow 25, nebulizer gas 1 (GS1) 15, heater gas 2 (GS2) 30 and interface heater at 150°C. The mass spectrometer acquired 250ms full scan TOF-MS data followed by up to 30, 50ms full scan product ion data, with a rolling collision energy, in an Information Dependent Acquisition (IDA) scan mode. Full scan TOFMS data were acquired over the mass range m/z 350-2000 and for product ion ms/ms, m/z 100-1500. Ions observed in the TOF-MS scan exceeding a threshold of 150 counts and a charge state of +2 to +5 were set to trigger the acquisition of product ion, ms/ms spectra of the resultant 30 most intense ions. The data were acquired and processed using Analyst v1.7 software (SCIEX). Protein identification was carried out using Protein Pilot 5.0 (SCIEX) for database searching.

### Proteomics Data and GO Analysis

A network analysis was carried out using the STRING protein query database for Mus musculus using the official gene-symbol (https://string-db.org) ^56^. The confidence score cut-off was set as 0.7. The MS data were analysed for GO terms enrichment using Enrichr ^57-59^ and the results for the top 10 most enriched terms are reported, with the length of each bar being proportional to -Log10 (*P*) where *P* represents the *P* value computed using the Fisher exact test.

### Single-molecule Localization Microscopy and Analysis

Primary cortical neurons were cultured and imaged on a glass bottom dish (Cellvis D29-20-1.5-N) as reported previously ^60^. Cells were then transduced with G3BP2-mEos3.2 and CIRTS and images were acquired on an ELYRA PSI microscope equipped with a x100/1.46 α Plan-Apochromat oil-immersion objective and an EMCCD camera. sptPALM was performed as reported by ^61^. Briefly, G3BP2-mEos3.2-expressing cells were photo-converted with a 405 nm laser and excited using a 561 nm laser. 16,000 frames were acquired at a rate of 33 Hz. All data were acquired using Meta-Morph Microscopy Automation and Image Analysis software, version 7.7.8 (Molecular Devices) and further processed using PalmTracer software ^61^.

### Statistical Analysis

Statistical analyses were performed using GraphPad Prism 9 unless otherwise specified. An unpaired two-tailed Student’s *t-*test was performed when comparing two categories. When more than two groups were compared, one-way ANOVA followed by a Dunnett’s multiple comparisons test were used. Results are mean (n ≥ 3) ± standard error of the mean (s.e.m.) unless otherwise stated.

### Data and Code Availability

All sequencing data and analysis pipeline are available on GitHub at https://github.com/Qiongyi/lncRNA_2020. Datasets generated during this study have been deposited and publicly available at http://genome.ucsc.edu/cgi-bin/hgTracks?db=mm10&hubUrl=https://data.cyverse.org/dav-anon/iplant/home/qiongyi/lncRNA2020/hub_lncRNA2020_v1.0.txt.

## Supplemental Figure Legend

**Supplemental Figure 1.**
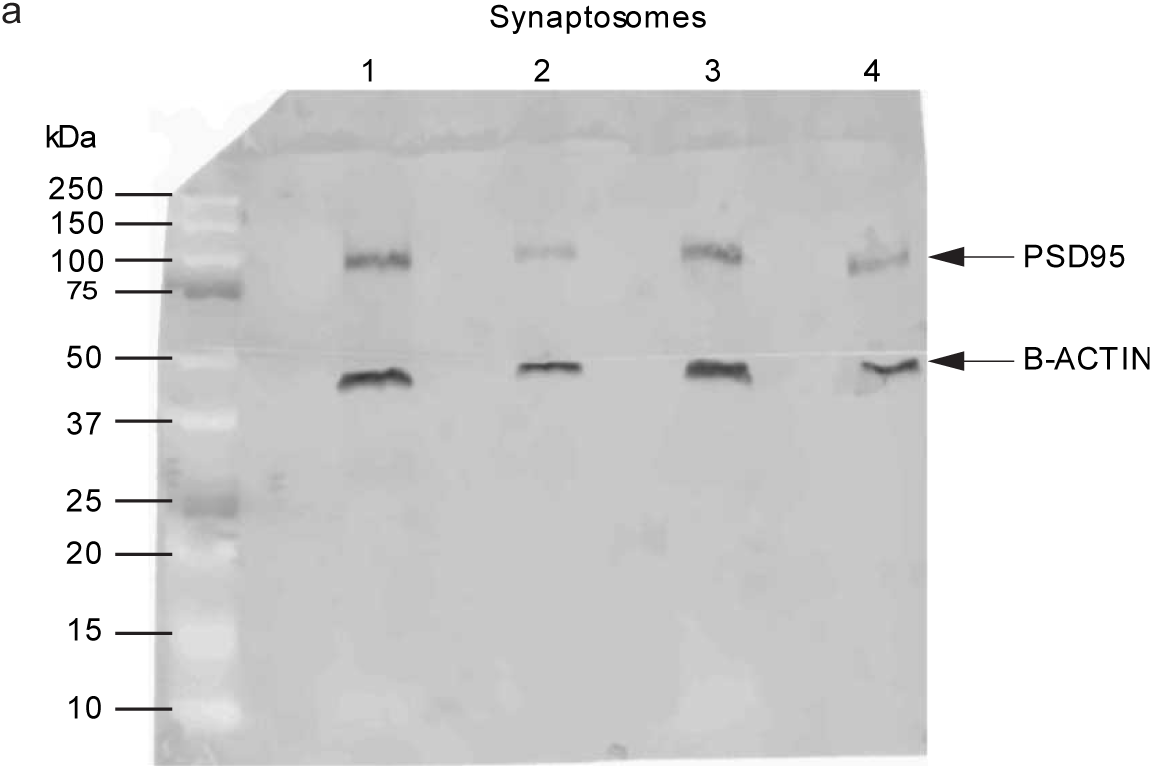
(A) Blotting of PSD95 protein after synaptosome isolation from different cohorts of mice. Each lanes represents separate synaptosome fractions. Synaptosome marker, PSD95 and the internal control are indicated.

**Supplemental Figure 2.**
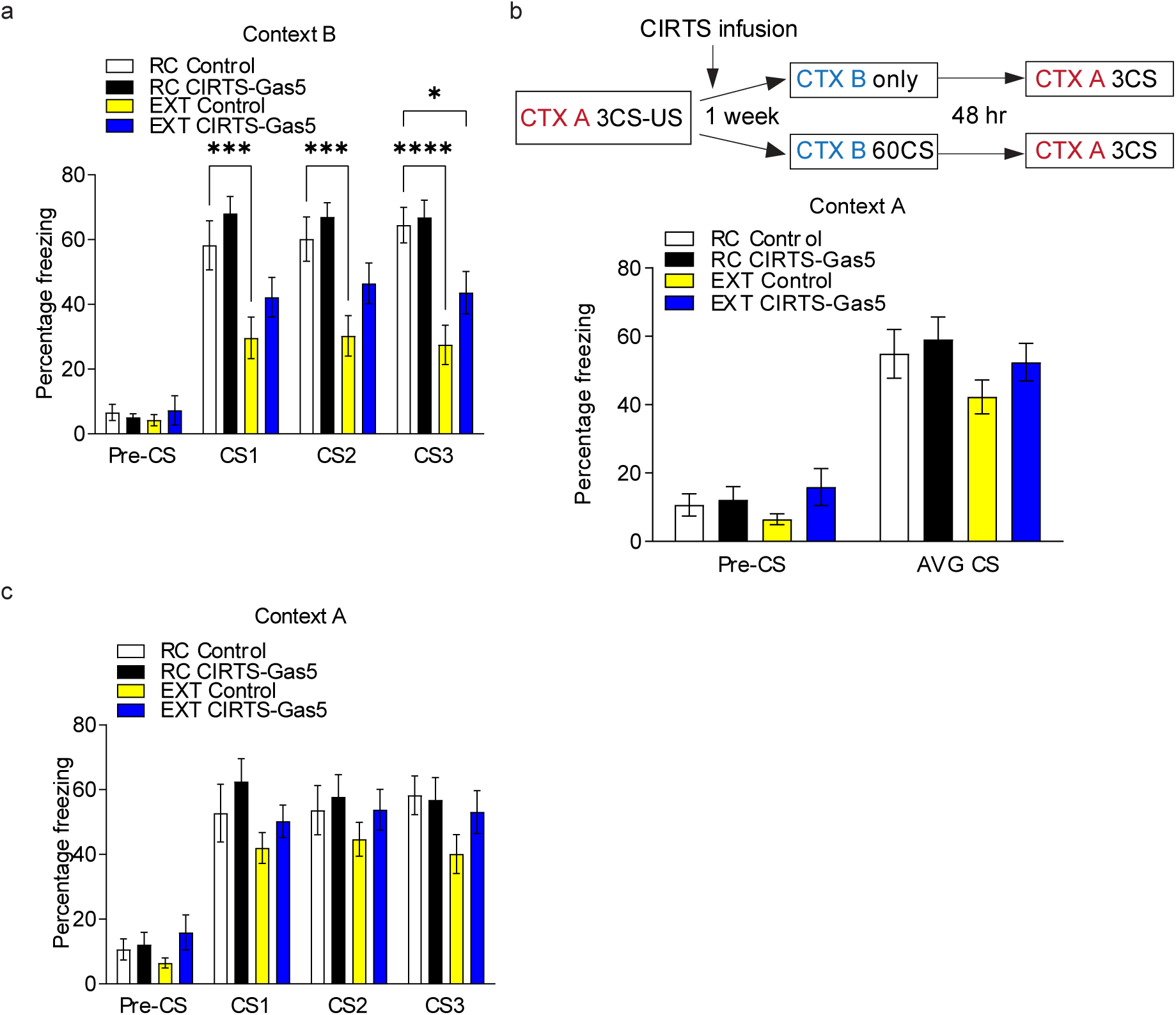
(A) Bar plot showing all 3CS in context B fear extinction test. Knockdown and behavioural condition are identical to Figure 3. n > 5 biologically independent animals per group, repeated two-way ANOVA with Dunnett’s posthoc tests. **** p <0.001; *** p<0.005; * p<0.05. (B) Top panel: Schematic of the behavioural protocol used to test the effect of Gas5 knockdown in the ILPFC on fear memory. CTX, context; CS, conditioned stimulus; US, unconditioned stimulus. Bottom panel: Bar plot showing context A 3CS test. Animals were subjected to similar conditions as in Figure 6f. (C) Bar Plot showing all 3CS in context A fear test. Knockdown and behavioural conditions are identical to Figure 6. n > 5 biologically independent animals per group, repeated two-way ANOVA with Dunnett’s posthoc tests.

