## Supplemental Table 2 for "Fear extinction is regulated by long noncoding RNA activity at the synapse"

**Supplemental Table S2**

**Primers for quantitative PCR**

| Name | Forward Sequence (5’-3’) | Reverse Sequence (5’-3’) |
|---|---|---|
| Exon 11 - Intron 11 - Exon 12 | GCTCCTGTGACAAGTGGA CA | GCACCTGCAACAACCAAC AG |
| Intron 9 - Exon 10 | CTGTGCCACTGACTTAAC CC | TCAGAATGAAGGACCGGA AA |
| Intron 8 - Exon11 | AGGTACTGTTAGTGATGA TC | ATGTCCACTTGTCACAGG AG |
| Exon 2 - Intron 2 | AATGGCAGTGTGGACCTC TG | TCAGTCCCCTTCTTCACGG A |
| Intron 5 | GAACAAATACTGACTACC TG | TGCCCCACAATAAATGTC AA |
| Exon 1-Exon 2-Exon 3 | ATTCTGAGCAGGAATGGC AG | TCCTCCTTTGCCACAGAAC T |
| Exon 8-Exon 9-Exon 10-Exon 11-Exon 12 | GTACAAATAATGGTTTGA AT | TGTTATAATACACTTTAAT G |
| *Gas5* | GGAAGCTGGATAACAGA GCGA | GCATGCAACCAGTTAACT TTCA |
| 18S rRNA | CTGGATACCGCAGCTAGG AA | GAATTTCACCTCTAGCGG CG |
| *Pgk* | TGCACGCTTCAAAAGCGC ACG | AAGTCCACCCTCATCACG ACCC |
| *Rpph1* | CTCTACGCTTGGGCAGAC | CTCACCTCAGCCATTGAA CT |
| *Rmrp* | CCTGTTTCCTAGGCTACA TACG | GCGGGCTAACAGTGACTT |
| *Rn7sk* | TCGGTCAAGGGTATACGA GTAG | TTTGGATGTGTCTGGAGTC TTG |
| *9330121K16Rik* | AGTAGAGTTAGGGTGGGA TAGG | ACAGGCACTGATGTGAGT TAG |
| *Gm28437* | CAGGATTCTTCTGAGCGT TCTAT | TGGGACTTCTAGAGGGTT AAGT |
| *GM47305* | AGACAGGAGGATCGCTTG A | TCACCATATTGATGCCGA ACTTA |
| *Cyrano* | GGCTCCATAGAAGCGACA TAC | CCCAAGAGCTGGGCATAA A |
| *Meg3* | ATTAGGCCAAAGCCATCA TCT | GGCGCTTCCAATCGATTT AC |

**Primers for *in-vitro* Transcription**

| Name | Sequence (5'-3') |
| --- | --- |
| T7 *Gas5* del1-50 | TAATACGACTCACTATAGGGTGATGGGACATCTTGTG |
| T7 *Gas5* | TAATACGACTCACTATAGGAGCCTTTCGGAGCTGTGC |
| *Gas5* Reverse | TTCATGTTATAATACACTTT |
| *Gas5* del458-504 Reverse | ATTTGAGCCTCCATCCAGGC |

**CIRTS gRNA**

| Name | Sequence (5'-3') |
| --- | --- |
| *Gas5* gRNA | AGCAAGCCAGCCAAATGAACAAGCATGCAA |
| Control Scramble | GATACATCATCTCTGTATTAGGCTCCCAAC |
